# Species interactions, divergence, and the rapid evolution of ecological sexual dimorphism in threespine sticklebacks

**DOI:** 10.1101/2025.10.26.684666

**Authors:** S. A. Blain, M. Roesti, K. A. Thompson, M. H. Kinney, D. Schluter

## Abstract

Variation in ecological sexual dimorphism (ESD), defined as differences between the sexes in ecologically-relevant traits, is a common feature of adaptive radiation, yet its causes remain unclear. Competition between the sexes for alternative resources can promote evolution of ESD when interspecific competition is reduced (competition hypothesis). Alternatively, sex-specific selection on ecological traits might weaken under strong directional selection in new environments (divergence hypothesis). We tested these hypotheses via their expected evolutionary outcomes in threespine stickleback populations from southwestern Canada. We found striking among-population variation in magnitude of ESD. Consistent with the divergence hypothesis, dimorphism along the main axis of body shape variation was reduced in recently derived freshwater populations compared to their contemporary marine ancestor. However, dimorphism only weakly declined with increasing phenotypic divergence. Average dimorphism along major freshwater body shape axes was similar between solitary populations and those coexisting with a competing species, contrary to the competition hypothesis. Instead, sympatry with a Benthic ecotype led to increased sexual size dimorphism in Limnetics, and total shape dimorphism was elevated in the sympatric stickleback species compared with solitary populations. In contrast to the mechanisms considered in existing theory, interactions in sympatry might produce elevated ESD by generating novel sex-specific selection pressures.

## Introduction

Variation among species in ecological sexual dimorphism (ESD), whereby the sexes of a species differ in ecologically-relevant traits, is a common feature of adaptive radiation (Butler et al., 2007; De Lisle & Rowe, 2017; Parsons et al., 2015; Wasiljew et al., 2021). One explanation for variation in ESD is that sex-specific selection weakens when populations diverge from an ancestral phenotype and adapt to new environments because both sexes experience similar directional selection, at least for a time (Connallon et al., 2018; De Lisle et al., 2018). An alternative explanation is variation in resource competition from other species (Bolnick & Doebeli, 2003; Li & Kokko, 2021; Stuart et al., 2021). In the absence of interspecific competition, depletion of shared resources between the sexes can lead to disruptive selection (Wilson & Turelli, 1986), which can favour alternative phenotypic extremes in males and females. Provided that genetic variation for resource-acquisition traits is at least partly independent between the sexes, greater differences between the sexes are expected to evolve (Bolnick & Doebeli, 2003; Cooper et al., 2011). Experimental work supports this mechanism: manipulating density and sex ratio produced disruptive selection on sexually dimorphic feeding morphology in salamanders, measured via individual growth (De Lisle & Rowe, 2015), and elevated ESD evolved in Drosophila under high relative to low competition (De Lisle, 2022). Evidence from nature, however, is scarce and mixed: comparative analyses across gradients of species richness report variable effects on ESD (Meiri et al., 2014; Pincheira-Donoso et al., 2018; Stephens & Wiens, 2009; Stuart et al., 2021).

Hereafter, we refer to these alternative explanations as the ‘divergence’ and the ‘competition’ hypotheses. Additional possibilities include variation in sex-specific selection caused by ecological factors that affect mate choice or reproductive success (De Lisle, 2019). A comparative approach that leverages variation in ESD among closely related populations or species in different environments provides an opportunity to test the alternatives (Stephens & Wiens, 2009; Stuart et al., 2021).

We evaluated the competition and divergence hypotheses using threespine stickleback (*Gasterosteus aculeatus* L.) from the ocean and from recently formed freshwater lakes in coastal British Columbia, Canada (Taylor & McPhail, 2000; Vamosi, 2003). To test the divergence hypothesis, we compared ESD in stickleback that recently colonized and adapted to freshwater environments with ESD in the ancestral marine environment (Bell & Foster, 1994). We further asked whether freshwater populations more divergent from the marine form have evolved an even greater reduction in sexual dimorphism than less divergent freshwater populations, as would be expected under greater net directional selection since colonization (Fig 1B). All of the freshwater populations are of a similar age and evolved independently after marine stickleback colonized these coastal lakes following deglaciation, roughly 12,000 years ago (Bell & Foster, 1994; Hohenlohe et al., 2010; Magalhaes et al., 2021; Morris et al., 2018; Roesti et al., 2025). The predictions primarily concern ESD along the axis of divergence between ancestral and derived environments, but we also consider the possibility of dimorphism evolution along alternative axes (Cooper et al., 2011). It has been previously observed that Benthics from Paxton Lake exhibit less sexual dimorphism in body shape than the Pacific marine stickleback (Albert et al., 2008), that stream populations with a benthic phenotype have reduced dimorphism relative to marine stickleback (Kitano et al., 2012), and that pelagic freshwater populations have relatively high levels of body shape dimorphism compared to less pelagic populations (Spoljaric & Reimchen, 2008). Here we tested whether ESD changes consistently in freshwater populations with increasing differentiation from marines.

**Fig. 1.**
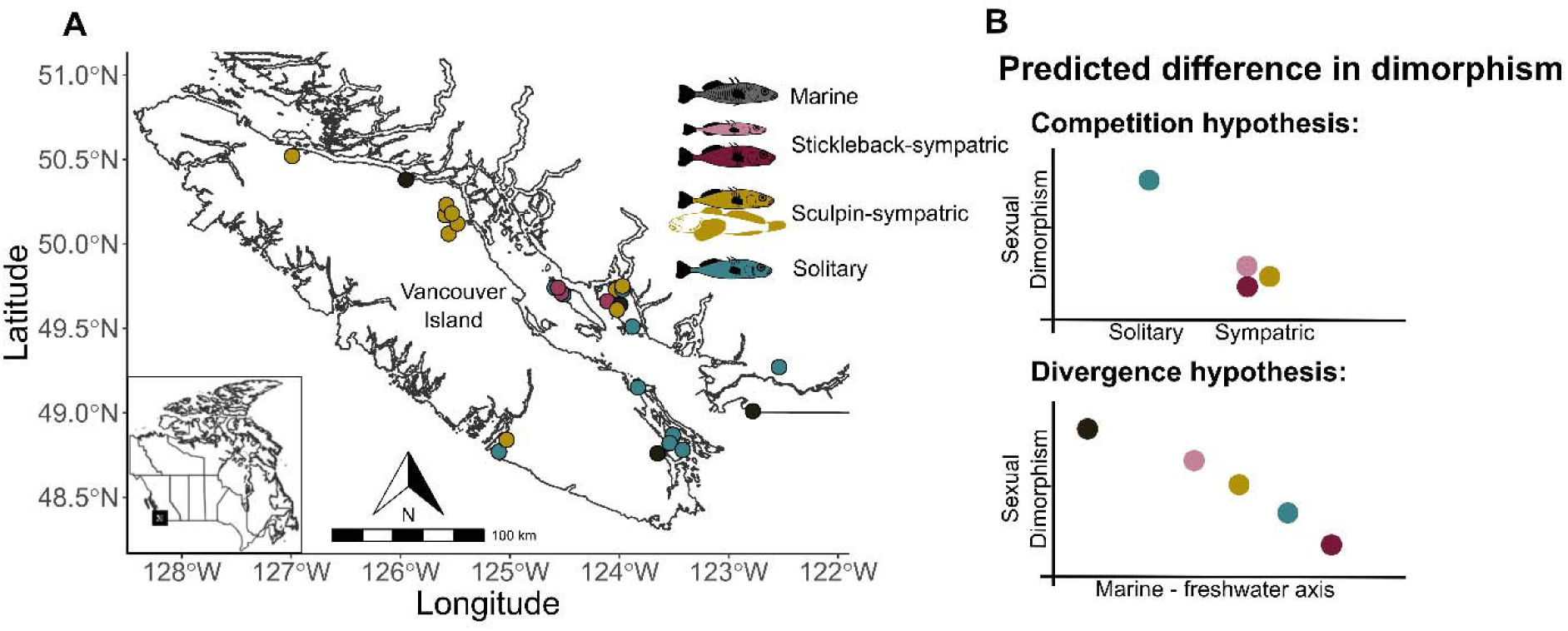
(A) Map of populations included in the study and population types. Stickleback-sympatric populations co-occur as species pairs, whereas sculpin-sympatric populations co-occur with the prickly sculpin. “Solitary” populations occur without a competitor fish population, and the marines represent the ancestral form. (B) Predicted levels of sexual dimorphism. Under the competition hypothesis, dimorphism is predicted to be higher in solitary populations than in stickleback-sympatric or sculpin-sympatric populations. Under the divergence hypothesis, dimorphism is predicted to be highest in marine stickleback and decrease with increasing divergence in freshwater populations.

Under the competition hypothesis, ecological sexual dimorphism should be greater when competing species are absent from lakes. We examined this prediction using two comparisons (Fig 1A, B; Table S1). First, several lakes contain a pair of reproductively isolated and ecologically divergent “Benthic” and “Limnetic” stickleback species. These sympatric ecotypes differ markedly in numerous traits associated with resource acquisition and habitat use, including gill raker number and length, overall body shape, head and jaw morphology, and bony defensive armor (McGee et al., 2013; Schluter & McPhail, 1992; Vamosi & Schluter, 2004). Their exaggerated divergence in sympatry is partly the evolved outcome of interspecific resource competition (Rundle et al., 2003; Schluter, 2003). We compared ESD in these sympatric stickleback ecotypes to “solitary” stickleback populations – generalist stickleback that occur alone in otherwise similar lakes and exhibit an intermediate phenotype (Nagel & Schluter, 1998; Schluter & McPhail, 1992). Second, we compared ESD in solitary stickleback populations with that in single-species stickleback populations sympatric with prickly sculpin (*Cottus asper* Richardson), a competitor and occasional intraguild predator (Roesti et al., 2023; Vamosi, 2003). These “sculpin-sympatric” stickleback populations typically have more gill rakers, a narrower gape, a more elongated body shape, enhanced defensive armor, as well as a more pelagic diet compared to solitary populations (Ingram et al., 2012; Miller et al., 2015; Roesti et al., 2023). Solitary populations have increased phenotypic variation and greater phenotypic plasticity in body shape compared with sympatric and marine populations, consistent with phenotypic evolution in response to intraspecific competition (Svanbäck & Schluter, 2012). The competition hypothesis mainly predicts stronger ESD along the same phenotypic axis as divergence between the species (Bolnick & Doebeli, 2003), but we also consider the possibility that ESD may have evolved along alternative axes (Cooper et al., 2011; De Lisle & Rowe, 2017).

## Methods

### Specimen sampling

We collected wild adult threespine stickleback between April and July in 2015 to 2018 using minnow traps and dip nets. Stickleback-sympatric populations (Benthics and Limnetics) were sampled from three lakes: Little Quarry, Paxton, and Priest (Fig. 1A; Table S1). Sculpin-sympatric stickleback populations were sampled from nine lakes: McNair, Roselle, Paq, Morton, Blackwater, Ormond, Merrill, North and Pachena lakes (Fig. 1A; Table S1). Solitary stickleback populations were sampled from ten lakes: Weston, Bullocks, Black, Kirk, Cranby, Maxwell, Mike, Trout, Klein, and Hoggan (Fig. 1A; Table S1). Cutthroat trout (*Oncorhynchus clarkii*), a predator of both stickleback and sculpin, have historically co-occurred with all freshwater stickleback populations (Roesti et al., 2023; Vamosi, 2003). We also sampled marine (anadromous) stickleback from the Cowichan and Little Campbell rivers, and saltwater marine fish from Oyster Lagoon (Fig. 1A; Table S1).

Because all lakes occur in different drainage basins (except Klein and North lakes, which are, however, geographically separated and are clearly genetically distinct; Miller et al. 2019), and lake types are geographically interspersed, we assume that freshwater populations evolved independently from marine ancestors following postglacial colonization (Miller et al., 2019; Taylor & McPhail, 2000). Independent, parallel adaptation to freshwater has been established in most, but not all, of these populations (Jones, Chan, et al., 2012; Miller et al., 2019; Roesti et al., 2023, 2025). All samples were euthanized in the field with buffered MS-222 and preserved in 95% ethanol. We targeted a sampled size of ten individuals per sex in each population (Table S1).

### Phenotyping

Specimens were gradually transitioned from 95% ethanol to water via serial dilutions over five days and then fixed in 10% formalin for at least 48 hours. Samples were then stained with alizarin red to enhance visualization of bony structures following standard procedures (Peichel et al., 2001). Sex was determined by the presence of testes or ovaries during dissection. Each specimen was photographed on the right side unless that side was visibly damaged.

We obtained *x-* and *y-*coordinates of 25 fixed landmarks from each specimen photograph using tpsDig (Rohlf, 2018). We used the same landmarks as in Albert et al. (Albert et al., 2008), except for the two pelvic girdle landmarks (4 & 5), which were omitted because the girdle was absent in most Benthics from Paxton and Little Quarry lakes (Fig. S1). These landmarks represent features associated with locomotion (fin insertions, body depth), defense (spine locations), and foraging (eye position and size, head size, snout and jaw shape) (Arnegard et al., 2014; Willacker et al., 2010). We performed a Procrustes analysis to scale, coordinate, and align landmarks across specimens using the *procGPA()* function in the *shapes* R package (Dryden, 2021) and used the resulting *x-* and *y-*coordinates of landmarks in subsequent analyses of shape variation. Average sexual shape differences between marine and freshwater ecotypes are illustrated in Fig. S2.

In addition to overall body shape, we measured the following defensive armor traits: length of the first and second dorsal spines, the pelvic spine, and the length of the pelvic girdle using calipers and counted lateral plates on the right side of each stained specimen. Missing armor traits were assigned a length of 0 because armor loss represents a genetically-based transition from marine to freshwater environments (Cresko et al., 2004; Jones, Grabherr, et al., 2012; Marchinko, 2009). We further measured the following foraging traits: number of gill rakers on the epibranchial and ceratobranchial of the main gill arch and the lengths of the three posterior-most gill rakers on the ceratobranchial using a dissecting microscope.

Linear traits (i.e. length traits, not including counts) were size-corrected using standardized major axis regression in the *smatr* R package (Warton et al., 2012) to account for variation related to age and growth rates among individuals of the same population and sex. Size correction via ordinary linear regression can generate bias when size ranges differ between the sexes of a population, as was the case within Limnetic populations, because of regression-to-the-mean effects when not all variation in measured size is causal (Warton et al 2006). Thus, we first regressed each trait on standard length combining all fish in one model, assuming equal slopes across sexes and populations. We then calculated the adjusted trait value of each fish by adding its residual to the predicted value for its group at the grand mean standard length. Armor measurements of 0 (indicating a missing spine or pelvic girdle) were removed from the data set before size correction and then added back afterward.

A composite metric of gill raker length was obtained using the first principal component from a principal components analysis (PCA) of the covariance matrix of the size-corrected lengths of three gill rakers. A composite armor metric was obtained using the first principal component from the correlation matrix of number of lateral plates, size-corrected length of pelvic girdle, and size-corrected lengths of first dorsal, second dorsal, and pelvic spines. We used a correlation matrix for armor because the traits were not measured on the same scale. We used the *nipalsPca()* function of the *pcaMethods* R package (Stacklies et al., 2007) because it can tolerate a small number of missing values.

We visualized outliers using histograms and scatter plots of landmark coordinates and linear traits. Landmark data are inherently noisy because preservation can deform specimens, distorting landmark placement and additionally contributing to roll, yaw, and pitch in digitized images, leading to high variation in individual measurements. Hence, individual measurements ≥ 4.5 standard deviations from the grand mean were set to missing before size correction and before calculation of the composite gill raker length and armor traits.

### Visualizing sexual dimorphism

Trait differences between males and females were visualized using paired-data plots of predicted means from linear mixed-effects models (see *Statistical analysis*). Sex, ecotype (marine plus the four freshwater ecotypes) and their interaction were fixed effects; population and the interaction between sex and population were random effects. This ‘ecotype model’ allows trait means and sexual dimorphism to vary freely among populations.

### Testing the divergence hypothesis

We tested the divergence hypothesis by comparing sexual dimorphism in overall body shape, gill raker traits, armor traits, and size between marine and freshwater populations, and among freshwater populations in relation to their position along the major axis of marine–freshwater shape variation. We calculated this shape axis with a PCA on the covariance matrix of body shape landmark coordinates using all individuals from marine and freshwater populations, using *nipalsPca()* (Stacklies et al., 2007). Landmarks were already scaled to the same centroid size and were measured in the same units. Hereafter, PC1_M-FW_ from this analysis, which explained 21.5% of the phenotypic variation in the 50 landmark traits, is referred to as the ‘marine – freshwater axis’ because marine fish were at one extreme and the most divergent freshwater populations were at the other. Among-population variance in PC1_M-FW_ accounted for 35.5 % of total among-population variance across landmarks.

Differences in sexual dimorphism between marine and freshwater populations along the marine – freshwater axis were tested using a linear mixed-effects model similar to the ecotype model (see *Visualizing sexual dimorphism*). Individual PC1_M-FW_ scores were the response variable, with sex, habitat (marine vs. freshwater), and their interaction as fixed effects, and population and population × sex as random effects. The interaction indicated whether sexual dimorphism differed between marine and freshwater populations.

The relationship between sexual dimorphism and position along the PC1_M-FW_ axis was estimated by fitting a mixed-effects ‘divergence model’ to trait values of individuals (including PC1_M-FW_ scores) and regressing against population mean PC1_M-FW,_ with sex, the interaction between sex and PC1_M-FW_, habitat, and the interaction between habitat and sex included as fixed effects. Population and the interaction between population and sex (population variation in the amount of sexual dimorphism) were random effects. The interaction between sex and population mean PC1_M-FW_ indicates whether dimorphism changes continuously along the marine–freshwater shape axis. The interaction between sex and habitat allows dimorphism to differ between marine and freshwater populations. The relationship between sexual dimorphism and the marine-freshwater axis was visualized using the differences between model fitted values of male and female population means. Similar analyses were conducted to investigate variation in sexual dimorphism in standard length, gill raker traits, and armor along the marine–freshwater phenotypic axis.

### Testing the competition hypothesis

To compare solitary versus stickleback-sympatric populations, we performed a PCA on the covariance matrix of body shape landmark *x-* and *y-* coordinates from solitary, Benthic, and Limnetic (but not sculpin-sympatric or marine) populations. Hereafter, the major axis from this analysis (PC1_Lim-Ben_), which explained 20.7% of phenotypic variation in the 50 traits, is referred to as the ‘Limnetic – Benthic axis’. Among population variance in this PC axis accounted for 28.9% of the total variance among the populations. We tested the competition hypothesis by fitting a linear mixed-effects ‘sympatry model’ to PC1_Lim-Ben_ scores and other traits of individuals with sex, sympatry (solitary vs sympatric), and the interaction between sex and sympatry as fixed effects. Population and the interaction between population and sex were random effects. The interaction between sex and sympatry tested for differences in sexual dimorphism between solitary and stickleback-sympatric (Limnetic and Benthic) populations. Sexual dimorphism was visualized as the difference between male and female model-predicted means per population. Similar analyses were conducted for gill raker traits, armor traits, and body size.

We next tested the prediction that sexual dimorphism should be greater in solitary than sculpin-sympatric populations. A PCA on the covariance matrix of body shape landmark coordinates including only solitary and sculpin-sympatric specimens yielded PC1_Sol-Sym_, explaining 19.7% of the phenotypic variation in the 50 traits. Hereafter, this axis is referred to as the ‘solitary – sculpin-sympatric axis’. Among-population variance in this PC accounted for 14.8 % of total among-population variance. The same sympatry model was applied, with sex, sympatry (solitary vs. sculpin-sympatric), and sex × sympatry as fixed effects, and population and population × sex as random effects. Sexual dimorphism was visualized using the difference between male and female model-predicted means in each population. Similar analyses were carried out on the other phenotypic traits (armor, gill rakers, and body size).

### Total dimorphism

Although the competition and divergence hypotheses related primarily to dimorphism along specific axes of variation, it is possible that sexual dimorphism in body shape is primarily along other axes of variation. Indeed, sexual dimorphism in body shape within ecotypes was not parallel with divergence among ecotypes (Fig S3). Therefore, we also compared total sexual dimorphism in shape of marine, solitary, and sympatric populations as follows. For each population, we first computed landmark coordinate means and standard errors (SE) for males and females. We then calculated the difference between males and females (sexual dimorphism) in each trait and population, and the squared SE of dimorphism as the sum of the squared SEs of the male and female means. Total dimorphism was calculated as the sum of the absolute values of dimorphism in all landmark coordinates. The squared SE for total dimorphism was calculated as the sum of the squared SEs of the individual coordinates, assuming approximate independence. The “*x*3” coordinate (Fig S1, anterior insertion of anal fin) was left out of the calculation because it evolved little between marine and freshwater populations and represented an outlier in comparisons of dimorphism between ecotypes (Fig S4).

### Statistical analysis

We fit linear mixed-effect models using the *lmer()* function from the *lme4* R package (Bates et al., 2015), or, when singular fits occurred, with the *blmer()* function from the *blme* R package (Chung et al., 2013). The *blme* package uses penalized likelihood to prevent point estimates of zero for the variances of the random effects and yields slightly more conservative significance levels for fixed effects. Population and the interaction of population and sex were included as random effects to allow means and sexual dimorphism to vary among populations. ANOVA tables for *lmer()* fits used Type II sums of squares with Satterthwaite’s degrees of freedom from the *lmerTest* package (Kuznetsova et al., 2017). ANOVA tables for *blmer()* fits were generated using the Type II *Anova()* method of the *car* R package with Wald *F*-tests and Kenward-Roger degrees of freedom (Fox & Weisberg, 2019). All analyses treated populations as the independent unit of replication, based on our assumption of phylogenetic independence of populations (see above) which has been established for most but not all populations included herein (Miller et al., 2019; Roesti et al., 2025; Taylor & McPhail, 2000). For the same reason, we treated Benthics and Limnetics as independent points even though they coexist as pairs within the same lakes. Mean differences in sexual dimorphism between ecotypes and other groups were estimated from fitted models as contrasts using the *emmeans* R package (Lenth et al., 2021). Differences in total dimorphism between ecotypes were tested by fitting linear mixed models to population values weighted by their squared SEs using the *rma.uni*() function with REML estimation in the *metafor* package (Viechtbauer, 2010), with ecotype as a fixed effect and population as a random effect.

### Diet analysis

We assessed sexual dimorphism in diet in freshwater populations to test whether phenotypic measures of ESD correspond to differences between the sexes in resource use. Stomach content data were compiled from Schluter and McPhail (1992) and Roesti et al. (2023). Schluter and McPhail (1992) collected samples in both spring (May to June 1988) and fall (September) from Paxton, Priest, Enos, and Cranby Lakes, while Roesti et al. (2023) collected samples in spring (May to July 2015 and 2016). Enos Lake, which contained sympatric Benthic and Limnetic ecotypes in 1988 before they collapsed into a hybrid swarm by 1997 (Taylor et al., 2006), was included as a sympatric example even though it was not part of the morphological analysis. We additionally collected and examined the stomach contents of specimens from Trout and Klein lakes in September 2019, following the same protocols used by Schluter and McPhail (1992) and Roesti et al. (2023). To minimize foraging and digestion effects within traps, we only kept fish from traps that had been set for two hours or less. Individuals were euthanized immediately after being removed from traps and preserved in 95% ethanol. Each prey item was assigned to a taxonomic category (ranging from genus to order) and classified as “littoral”, “pelagic”, or “other” based on their taxonomy, following Schluter and McPhail (1992). For each fish, we then calculated the proportion of prey items belonging to each of those categories.

For each population, we calculated proportional similarity of males and female diets as 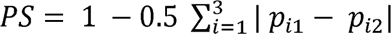 where *p*_*i*1_ and *p*_*i*2_ are the proportional contributions of prey category *i* in each sex (Bolnick et al., 2002). We calculated sexual dimorphism in diet as 1-*PS*. For populations that were collected in both spring and fall, dimorphism was estimated separately for each season. We had diet data from both spring 1988 and 2023 from Cranby lake, so we used the average of the sexual dimorphism estimates from the two samples. For each season, we evaluated the effect of sympatry by fitting linear models with population type (stickleback-sympatric, sculpin-sympatric, or solitary) as a predictor. We then evaluated whether sexual dimorphism was related to position along the marine – freshwater axis by fitting linear models for each season with PC1_M-FW_ as a predictor.

## Results

### Decline of sexual dimorphism in freshwater

The divergence hypothesis of sexual dimorphism was partly supported by patterns of body shape evolution. Freshwater stickleback repeatedly evolved reduced sexual dimorphism in body shape relative to the ancestral marine form (Fig. 2A). While all populations tended to be sexually dimorphic in shape, dimorphism in marine populations along the PC1_M-FW_ axis (82.1 + 12.0 SE) was more than twice that of freshwater populations (36.7 + 4.5 SE; Fig. 2B; sex × PC1_M-FW_ interaction: *F*_1,31.3_ = 12.62, *P* = 0.0012). Males were more freshwater-like along this phenotypic axis than females (Fig. 2A). The most sexually dimorphic landmarks in marine fish were those associated with snout length and overall head size, which were reduced in derived freshwater populations (Fig. S2). The only exception was the anterior insertion point of the anal fin (landmark #3, Figs S1, S2), whose *x*-coordinate was strongly dimorphic in both marine and freshwater populations.

**Fig. 2.**
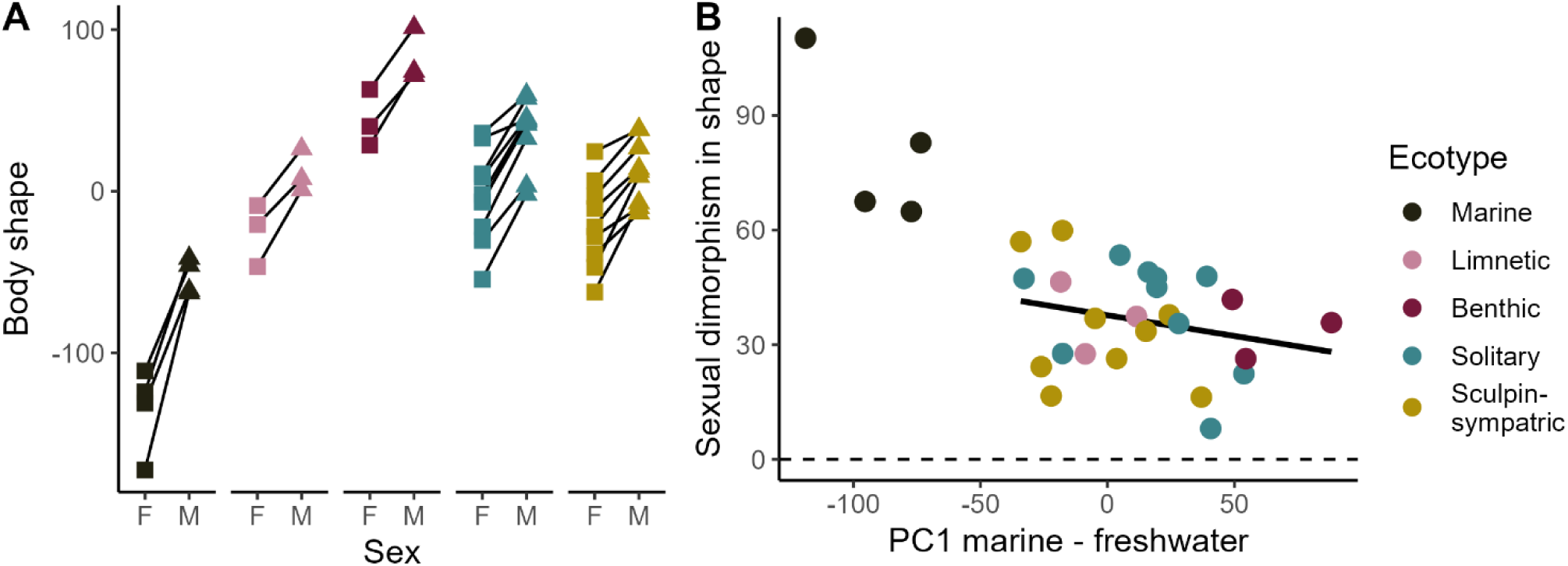
Variation in sexual dimorphism in body shape along the marine – freshwater shape axis PC1_M-FW_. (A) Female (F) and male (M) shapes for all populations and ecotypes. Lines connect model-predicted males and female means from the same population; steeper slopes indicate higher sexual dimorphism. (B) Sexual dimorphism in body shape in relation to population mean position on the principal marine – freshwater phenotypic axis PC1_M-FW_. Each point represents a single population, with dimorphism calculated as the difference between model-predicted male and female means.

In line with the second prediction of the divergence hypothesis, the magnitude of sexual dimorphism in body shape along the marine-freshwater axis also tended to decline in derived freshwater populations with increasing shape divergence from the ancestral marine form (Fig. 2B). However, the slope of this relationship was not detectably different from zero (divergence model: *F*_1,23.5_ = 0.38, *P* = 0.54). Total shape dimorphism was also greatest in marines (Fig. 3), yet variation among freshwater populations was not associated with increasing shape divergence from the ancestral marine form (linear model weighted by squared SE of total dimorphism; *F* = 0.029, *P* = 0.87).

**Fig. 3.**
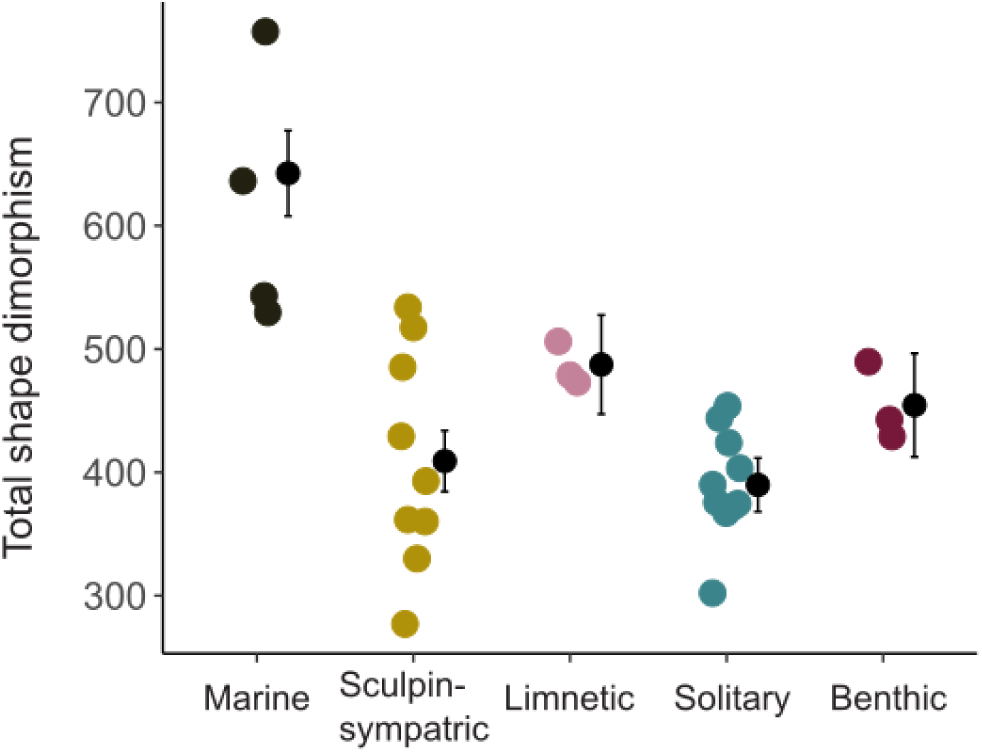
Total shape dimorphism in populations of each ecotype. Model-predicted values were obtained by fitting a linear mixed model to population values of total dimorphism weighted by their squared SEs, with ecotype as a fixed effect and population as a random effect. Total dimorphism is the sum of the absolute value of male-female differences in each trait, leaving out the *x*3 coordinate.

Greater sexual dimorphism in marines compared to freshwater stickleback was not detected in any trait other than body shape. Of the other traits tested, only gill raker length was sexually dimorphic, averaged over all populations (*F*_1, 25.1_ = 28.7, *P* < 0.001), with male stickleback having longer gill rakers than females (Fig. 4). However, sexual dimorphism in this trait was not significantly higher in marine than freshwater populations (*F*_1, 32.0_ = 0.21, *P* = 0.65) and showed no clear relationship with population mean position along the marine-freshwater shape axis (Fig. S5; *F*_1,24.5_ = 1.23, *P* = 0.28). Neither gill raker number (*F*_1, 35.2_ = 0.16, *P* = 0.69), armor (*F*_1,35.0_ = 1.14, *P* = 0.29), nor body size (*F*_1,33.6_ = 1.78, *P* = 0.19) showed a detectable difference in sexual dimorphism between marine and freshwater populations. Sexual dimorphism in these traits also showed no association with mean population position along the marine–freshwater axis (Figs. S 6-8; gill raker number: *F*_1,26.0_ = 0.009, *P* = 0.93; armor: *F*_1,27.5_ = 0.27, *P* = 0.61; body size: *F*_1,26.9_ = 0.77, *P* = 0.39). Nevertheless, sexual dimorphism in body size and possibly armor varied among ecotypes (depending on size correction; see below).

**Fig. 4.**
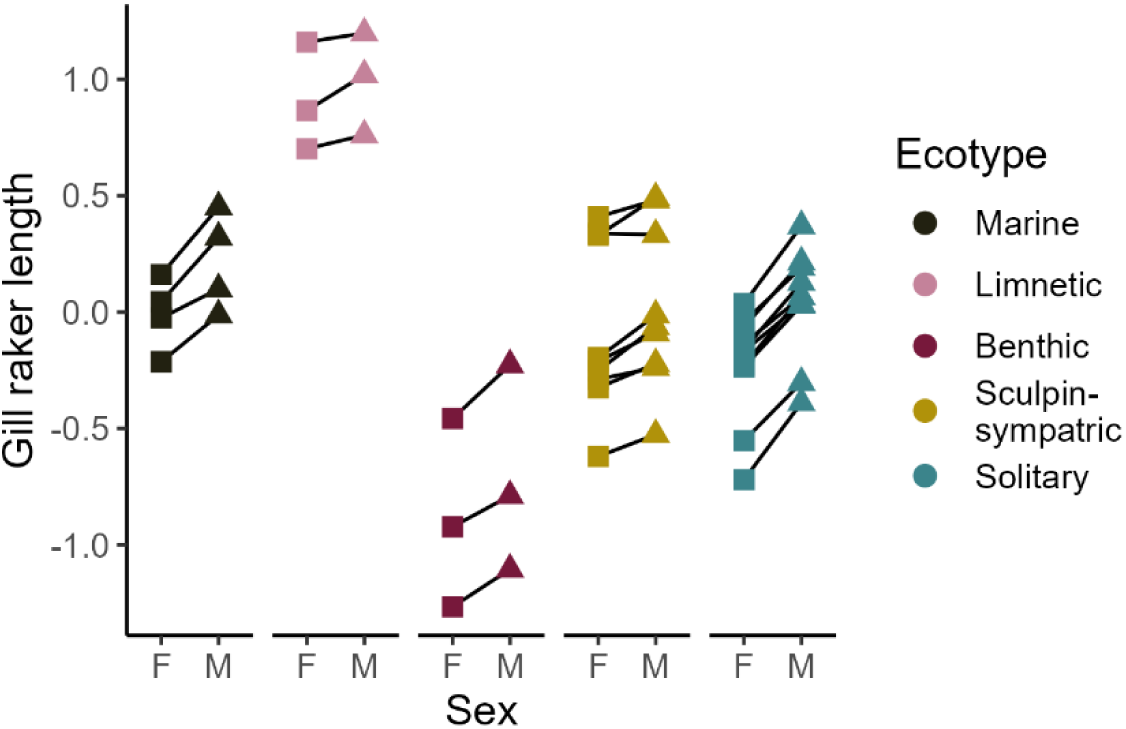
Sexual dimorphism in gill raker length. Lines connect model-predicted male and female means from the same population. Gill raker length is the principal axis of size-adjusted length traits.

### Sympatry and sexual dimorphism

None of the predictions of the competition hypothesis of ecological sexual dimorphism were upheld. Levels of body shape dimorphism along the Limnetic – Benthic axis were not noticeably greater in solitary than in stickleback-sympatric populations (Benthics and Limnetics combined) (Fig. 5A, Fig. S9; sympatry model: *F*_1,13.4_ = 1.94, *P* = 0.19). The same was true for gill raker number (Fig. S10; *F*_1,13.9_ = 2.47, *P* = 0.14) and the composite gill raker length trait (Fig. 4; *F*_1,13.4_ = 2.28, *P* = 0.15). Total sexual shape dimorphism was significantly greater in the sympatric than in the solitary populations (Fig. 3; 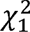 = 9.43, P = 0.002), opposite to the prediction.

**Fig. 5.**
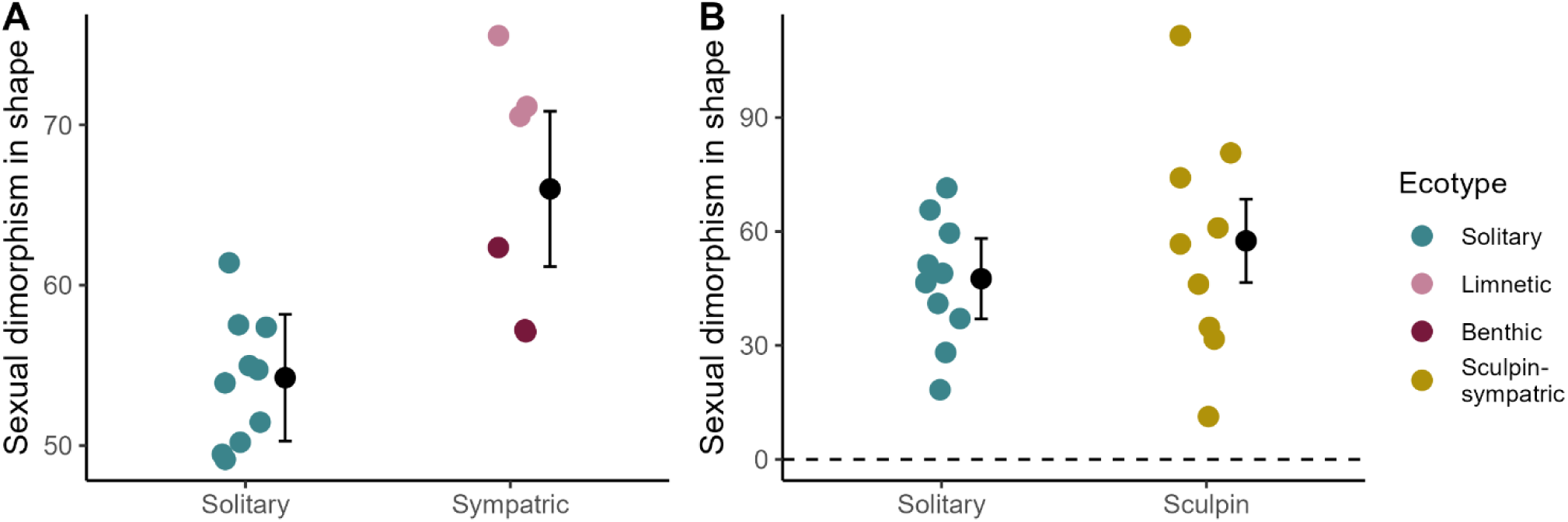
Sexual dimorphism in body shape in solitary stickleback versus that in stickleback populations sympatric with a competitor species. (A) Dimorphism in solitary stickleback vs members of stickleback species pairs. Each point represents sexual dimorphism in one population, the difference between male and female model-predicted means along the principal Limnetic – Benthic body shape axis PC1_Lim-Ben_. (B) Dimorphism in solitary vs sculpin-sympatric populations. Each point represents sexual dimorphism in one population, quantified as the difference between male and female model-predicted means along the principal solitary - sculpin-sympatric body shape axis PC1_Sol-Sym_. Black dots represent model-fitted mean dimorphism of a given population type, with standard error bars.

The direction of armor sexual dimorphism was significantly reversed in stickleback-sympatric populations compared to solitary populations (*F*_1,13.6_ = 5.38, *P* = 0.036), although the absolute value of dimorphism was not different between sympatric and solitary population (*F*_1,13.4_ = 0.58, *P* = 0.46). This reversal resulted mainly from enhanced armor in Limnetic females relative to body size (Fig. S11). Although Limnetic females are smaller (see below) and have less armor than males, they showed higher relative armor values after size correction, leading to reversed average sexual dimorphism in stickleback-sympatric relative to solitary populations (Fig. S11).

Similarly, levels of shape dimorphism along the Solitary – Sculpin axis were not detectably different between sculpin-sympatric and solitary populations (Fig. 5B, Fig. S12; *F*_1,17.2_ = 0.41, *P* = 0.53). The same was true for gill raker number (Fig. S10; *F*_1,15.9_ = 0.037, *P* = 0.85), armor (Fig. S11; *F*_1,18.6_ = 1.78, *P* = 0.20), and – marginally – the gill raker length trait (Fig. 4; *F*_1,16.6_ = 4.27, *P* = 0.055). Total sexual dimorphism in shape was also not significantly different between solitary and sculpin-sympatric populations (Fig. 3; 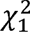 = 0.21, P = 0.65).

### Gain in sexual size dimorphism

In addition to greater total shape dimorphism in sympatric Limnetic and Benthic stickleback compared with solitary populations (Fig. 3), a striking pattern was the repeated gain of a novel pattern of sexual size dimorphism in one of the two sympatric stickleback species compared to all other ecotypes (Fig. 6). Whereas females were slightly larger on average in most stickleback populations, male Limnetics were – across all replicate stickleback-sympatric lakes – much larger than females (ecotype model: *F*_4,25.1_ = 8.78, *P* < 0.001). Sexual dimorphism in body size was also significantly greater in Limnetics than Benthics (ecotype model retaining only these two species: *F*_1, 4.2_ = 10.19, *P* = 0.03). This shift was observed only in stickleback-sympatric populations and not in sculpin-sympatric populations, suggesting that the presence of a sympatric stickleback species, rather than any generic benthic competitor, is responsible. In contrast, marine females were larger than males (Fig. 6), indicating that the direction of size dimorphism in Limnetics is derived, not ancestral.

**Fig. 6.**
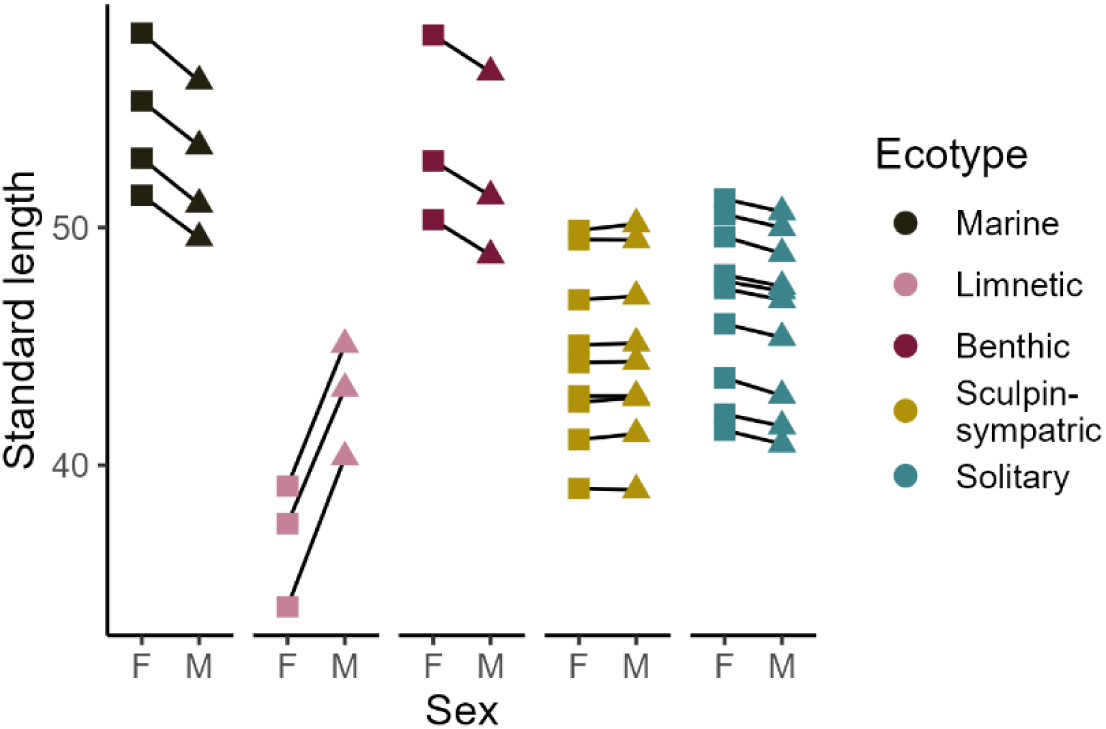
Variation in sexual size dimorphism (standard length, in mm). Each point represents the model-fitted mean standard length estimate for males or females from a population. Lines connect estimates for sexes from the same population.

### Sexual dimorphism in diet

Consistent with the divergence hypothesis, sexual dimorphism in diet declined in freshwater populations with increasing position along the marine-freshwater body shape axis. This pattern was most apparent in the spring diet samples (Fig. S13; PC1_M-FW_: *F*_1,17_ = 6.82, *P* = 0.02). The decline was not clearly detectable in the fall, but the sample size for this season was small (Fig. S13; PC1_M-FW_: *F*_1,5_ = 0.40, *P* = 0.56).

Again, in contrast to the prediction from the competition hypothesis, stickleback-sympatric and solitary populations exhibited similar levels of dimorphism in diet in both the spring and fall (Fig. S14; *F*_1, 12_ = 0.06, *P* = 0.82, *F*_1, 7_ = 3.5, *P* = 0.10). Interestingly, in the solitary populations – which have a fall diet intermediate between Benthics and Limnetics (Schluter & McPhail, 1992) – males consistently consumed more pelagic prey than females, while females consumed more prey items in the “other” category. This is consistent with longer gill rakers in males than females (Fig. 4), but is surprising because in body shape males are more similar to Benthics whereas females are more similar to Limnetics (Fig. 2). Likewise, levels of diet dimorphism were not detectably different between sculpin-sympatric and solitary populations in spring (*F*_1, 13_ = 1.28, *P* = 0.28).

## Discussion

Ecological sexual dimorphism (ESD) has evolved rapidly in threespine stickleback since freshwater populations formed at the end of the last ice age. Populations of the same ecotype showed relatively consistent levels of ESD, indicating that sex-specific trait divergence has evolved repeatedly and predictably within similar environments. We tested alternative explanations to identify the mechanisms underlying this variation, focusing on whether ESD reflects adaptation to new environments (the ‘divergence hypothesis’) or to shifts in interspecific competition (the ‘competition hypothesis’).

### Support for the divergence rather than the competition hypothesis

Under the divergence hypothesis, sexual dimorphism is predicted to decrease when both sexes experience similar directional selection during adaptation to new environments (Connallon et al., 2018). In alignment with this prediction, ESD in body shape was repeatedly reduced in derived freshwater compared to ancestral marine stickleback populations and sexual dimorphism in diet was reduced in more divergent freshwater populations. However, the second prediction of the divergence hypothesis, that is, a negative relationship between sexual dimorphism and the degree of phenotypic divergence from the marine ancestor, was only weakly supported. An alternative explanation, the competition hypothesis, predicts that solitary populations should evolve greater ESD than sympatric populations in a process analogous to ecological character displacement (Bolnick & Doebeli, 2003; De Lisle & Rowe, 2015; Li & Kokko, 2021). We tested this prediction by comparing solitary stickleback populations to sympatric stickleback species pairs and to stickleback populations that occur with prickly sculpin, a competitor for benthic resources. The prediction was not upheld in either case: levels of sexual dimorphism in body shape along the Benthic – Limnetic axis and along the sculpin sympatric – solitary axis were not greater in freshwater solitary populations compared to freshwater populations sympatric with other species. The same was true of gill raker number and length, armor, and diet. Instead, in contrast to all expectations, sympatric Limnetic stickleback exhibited higher levels of dimorphism in body size than Benthic or solitary stickleback populations, and both Benthics and Limnetics exhibited more total sexual shape dimorphism than solitary populations. Solitary and sculpin-sympatric populations had the lowest mean levels of total shape ESD.

Overall, these results were more consistent with the predictions from the divergence hypothesis, but only for shape and not for other morphological traits, while providing little empirical support for the competition hypothesis. The repeated reduction in body-shape dimorphism among freshwater populations likely reflects strong directional selection acting similarly on both sexes during the early stages of freshwater adaptation (Connallon et al., 2018). Ecological sexual dimorphism has been hypothesized to be a source of genetic variation that facilitated the rapid adaptation of ancestral marine stickleback to freshwater environments (Aguirre et al., 2008; Albert et al., 2008). However, if this hypothesis fully explained the observed pattern, we would expect the greatest reduction in dimorphism in the most phenotypically divergent freshwater populations. Although such a trend was observed in body shape along the marine – freshwater axis, it was not statistically significant and no trend was observed in total shape dimorphism.

Our finding that ESD is not greater in solitary populations aligns with most empirical results to date from other natural systems. For example, while sexual size dimorphism declines with species richness in assemblages of *Liolaemus* lizards (Pincheira-Donoso et al., 2018), similar levels of ecological sexual dimorphism are observed across communities of emydid turtles that vary in the number of competitor species present (Stephens & Wiens, 2009). Likewise, solitary *Anolis carolinensis* populations show similar levels of sexual dimorphism to those that co-occur with *A. sagrei*, a competitor species (Stuart et al., 2021), despite rapid ecological character displacement between the competitor species in sympatry (Stuart et al., 2014). Thus, while ecological character displacement between species evolves readily (Pfennig & Pfennig, 2010; Schluter, 2000a), analogous intersexual ecological character displacement within species appears uncommon in nature.

One explanation for the lack of ecological character displacement between the sexes is that competition-driven divergent selection is too weak or inconsistent between the sexes to drive repeated evolution across populations. Selection on dimorphic traits may fluctuate between the breeding season and other times of the year or between years (Reimchen & Nosil, 2004). Moreover, the magnitude of phenotypic differences between sexes is generally smaller than between species. Divergent selection is expected to be strongest at an intermediate levels of phenotypic similarity between competing populations (Schluter, 2000b), and the sexes may be too similar for strong intersexual divergent selection to arise. A second explanation is that solitary threespine stickleback populations may have evolved in other ways in response to intraspecific resource competition. Solitary populations exhibit wider niche breadths and greater phenotypic plasticity than both stickleback-sympatric and sculpin-sympatric populations (Roesti et al., 2023; Svanbäck & Schluter, 2012). These features are consistent with adaptation to resource competition within populations, but they need not result in sex-specific niche partitioning. The evolution of ecological sexual dimorphism may also be constrained by the high genetic correlation between sexes (Lande, 1980).

It is still possible that ecological sexual dimorphism is shaped by competition in stickleback. De Lisle (2019) suggested that threespine stickleback constitute a compelling case for the evolution of sexual dimorphism by ecological intersexual character displacement because dimorphism in ecologically-relevant traits has been observed repeatedly in stickleback, across both marine and freshwater populations (Aguirre & Akinpelu, 2010; Cooper et al., 2011; Kitano et al., 2007; McGee & Wainwright, 2013). Consistent with dimorphism evolving in response to competition, disruptive selection is reduced with higher levels of dimorphism across wild stickleback populations co-occurring with variable fish communities (Bolnick & Lau, 2008). However, selection favoring dimorphism may not differ systematically across competitive contexts. Although lakes containing solitary and competitor-sympatric populations overlap broadly in their physical and chemical characteristics, some (unmeasured) environmental variation could constrain both the co-occurrence of sympatric stickleback species and the evolution of ecological sexual dimorphism (Miller et al., 2019; Ormond et al., 2011; Vamosi, 2003). Therefore, the assumption that resource gradients are the same across lakes with and without competitor species may not fully hold (Bolnick & Doebeli, 2003; Li & Kokko, 2021). Moreover, sexual dimorphism in response to resource competition between the sexes is most likely to evolve with high individual specialization (Bolnick & Doebeli, 2003; Cooper et al., 2011), yet this is a feature of all freshwater populations and is unlikely to be greater in solitary relative to competitor-sympatric populations (Bolnick & Ballare, 2020; Matthews et al., 2010). Furthermore, if populations vary in the genetic architecture underlying sexual dimorphism, this could produce among-population differences independent of ecological context (Bolnick & Doebeli, 2003).

### Further explanations for variation in sexual dimorphism

Contrary to predictions from both hypotheses, Limnetic stickleback ecotypes from stickleback-sympatric lakes evolved elevated body size dimorphism relative to their marine ancestor, and both Limnetic and Benthic species showed consistently greater total shape dimorphism than solitary populations. One possible explanation is that ESD in these traits has evolved as a secondary consequence of ecological character displacement between the sympatric species (Schluter & McPhail, 1992). Perhaps the alternative lake habitats exploited by the two co-occurring species impose different selection landscapes for sexual dimorphism. Spoljaric and Reimchen (2008) reported that in their survey, the most sexually dimorphic stickleback populations were either marine or in lakes with a large pelagic zone, whereas the least dimorphic populations were in lakes with a proportionally larger benthic habitat. Similarly, Kitano et al. (2012) found reduced sexual dimorphism in body depth in benthic-like stream populations compared to marine populations. On the other hand, these trends would also be consistent with the divergence hypothesis.

Another possible explanation for elevated dimorphism in the sympatric species pairs is that they evolved via non-consumptive interactions. The prediction that ecological sexual dimorphism should decrease with species diversity assumes that the addition of species to the community generates no new sex-specific selection pressures via other interactions among more species – a condition that is unlikely to hold in stickleback, and likely many other organisms. For example, in the breeding season, male Limnetic stickleback build nests, court females, and care for eggs in the littoral zone of lakes and there they interact aggressively with males of the larger-bodied Benthics also defending territories (Larson, 1976). This interspecific male-male competition might generate selection for larger size in Limnetic males, and possibly larger head size in males of both species. In contrast, female Limnetics continue to forage primarily in the open water zone until egg-laying, where their small size confers a foraging advantage (Schluter, 1993). This hypothesis also suggests that sexual dimorphism in an ecologically-relevant trait can evolve as a by-product of adaptation to the different reproductive roles of males and females (Hedrick & Temeles, 1989; Nylin et al., 1993; Shine, 1989, 1991) that themselves might evolve in response to interactions between species. This mechanism seems most plausible in systems where traits that are involved in reproductive roles are also involved in resource acquisition, as in mouth-brooding cichlid fishes (Ronco et al., 2019).

Together, it is thus reasonable to expect that reproductive, sexual, and ecological selection interact to produce population-level variation in sexual dimorphism that does not conform to predictions from simple theory based on resource competition or directional selection. Quantifying selection on sexually dimorphic, ecologically-relevant traits both during and outside the breeding season will likely be key to disentangling these effects.

### Genetic basis of sexual dimorphism

Because all specimens were wild-caught, our data cannot fully disentangle the relative impacts of genetics and plastic sources in producing among-population variation in sexual dimorphism. Although males and females sampled from the same lake might be considered to be from a common environment, environmental conditions differ both between freshwater and marine environments and among lakes (Miller et al., 2019; Vamosi, 2003), potentially generating different plastic responses to sex-specific selection. In addition, the observed body size differences could emerge if male Limnetics had a longer lifespan than females, but little is known about lifespan variation of Benthic and Limnetic ecotypes. However, several lines of evidence suggest that at least some of the observed variation in ESD reflects evolved, rather than plastic, differences. Populations of the same ecotype exhibited similar magnitudes and patterns of ESD despite independent origins and geographic interspersion. Also, marine stickleback, which we found to exhibit greatest sexual dimorphism, have lower plasticity in shape than solitary freshwater sticklebacks, and similar plasticity to stickleback-sympatric populations (Svanbäck & Schluter, 2012), suggesting that greater dimorphism in marines is not due to greater overall phenotypic plasticity. Previous laboratory common garden experiments, including many of our study populations, have also confirmed a genetic basis of population differences in shape and other traits (Chhina et al., 2022; Day et al., 1994; Hatfield, 1997; Miller et al., 2015). Finally, we here reanalyzed shape variation in sexually mature F2 crosses between two Benthic and Limnetic population pairs (Conte et al., 2015), revealed QTL with sex-specific effects, confirming genetic variation for ESD (Supplementary Methods; Table S2). Establishing the genetic basis for variation in ESD across populations would help to better elucidate how this variation evolved.

### Conclusion

Overall, our study highlights the rapid evolution of ecological sexual dimorphism in threespine stickleback populations associated with adaptation to new environments. Although sexual dimorphism among derived freshwater populations was not clearly related to the magnitude of divergence from the ancestral marine form, sexual dimorphism in body shape was consistently reduced in freshwater relative to marine stickleback, thus aligning with the divergence hypothesis. On the other hand, the lack of increase in sexual dimorphism between solitary populations and those sympatric with a competitor species challenges the competition hypothesis and is in line with recent results from other organisms. Nevertheless, the repeated emergence of strong sexual size dimorphism in Limnetic stickleback co-occurring with Benthic stickleback within the same lakes, and the elevated levels of total dimorphism in both these sympatric ecotypes compared to the marine ancestor, underscore the role of interspecific interactions as drivers of sex-specific selection. Such interactions may represent a more potent source of variation in ecological sexual dimorphism within adaptive radiations than is predicted by existing ecological theory.

## Data accessibility

All data used in this study, including phenotypic trait data and stomach content data, have been deposited on Borealis. Code for processing and analyzing these data have also been deposited in the same repository. An anonymous reviewer version of this dataset is available here: https://borealisdata.ca/privateurl.xhtml?token=42ff349d-0552-4e37-813a-4f901203557c

