## Supplementary Materials for "Species interactions, divergence, and the rapid evolution of ecological sexual dimorphism in threespine sticklebacks"

**Supplementary Methods**

We reanalyzed data from Conte et al (2015) to test for the presence of QTL underlying differences between Benthics and Limnetics in ecological sexual dimorphism. Conte et al generated an F1 hybrid cross between a Benthic male and a Limnetic female from each of Priest and Paxton lakes in 2009 and reared them in the laboratory. In 2010 35 F1 hybrid adults (19 female and 16 male) from the Paxton cross and 25 F1 hybrid adults (12 female and 13 male) from the Priest cross were introduced two separate experimental ponds on UBC campus (described in Arnegard et al., 2014). Sexually mature F2 individuals were sampled in the spring of 2011, euthanized, stained and measured using the same landmarks as those used herein.

407 F2 fish from the Paxton Lake cross and 324 F2 fish from the Priest Lake cross were genotyped along with all parental individuals at hundreds of markers using a SNP array, which was used to determine parentage, phase, and to generate linkage maps (Conte et al., 2015). We imported the landmark measurements and genotype data into the R *qtl* package assuming an F2 intercross design (Broman et al., 2003). We used the *fill.geno*() command to impute missing genotypes and then converted values to a numeric score representing the fraction of each genotype inherited from the limnetic grandparent (0 if both alleles were benthic, 1 if both were limnetic, and 0.5 if a heterozygote). We used only autosomal markers to map sexually dimorphic traits.

To investigate the genetics of ecological sexual dimorphism we focused on landmark coordinates of the head and jaw, which show the greatest variation among ecotypes, especially the *x*-coordinates determining overall variation in head and snout length (Fig. S2, Fig. S4). In each cross, LDx was calculated as the first discriminant function separating the sexes in antero-posterior position of landmarks of the head region, including the trait “x4” (the posterior most point of the ectocoracoid) and all anterior $x$-coordinates (Fig S1). We also included LDxy, calculated as the first discriminant function separating the sexes in both $x$- and $y$- coordinates of landmark 4 (the posterior most point of the ectocoracoid) and all anterior traits (Fig S1).

Traits were mapped by regressing each composite trait simultaneously against all marker genotypes using the *glmnet* package in R (Tay et al., 2023). We set $\alpha = 1$, employing lasso regression with a likelihood penalty to prevent overfitting when modeling many possible explanatory variables such as marker genotypes (Tibshirani, 1996). F1xF1 family identity was included as a covariate in all analyses. We fit three models to the data from each lake. The first included only sex and family and represents the baseline ("covariate model"). The second model added the marker genotypes as main effects ("main effects model"). A significant main effect indicates segregating autosomal genetic variation for shape in both sexes. Detected markers are linked to autosomal genes that cause both males and females to vary along the axis of dimorphism in head shape. The third model included interactions between sex and genotype ("ESD model"), detecting the presence of autosomal genetic variation having greater effects in one sex than the other. These differential effects identify a genetic basis for the evolution of differences between populations in the amount of ecological sexual dimorphism. The regression parameter $\lambda$, which controls the magnitude of the penalty, was estimated when fitting the main effects model by minimizing the cross-validation score using the *cv.glmnet*() function. The process is stochastic and so was repeated 50 times, and we used the average of the 50 $\lambda$ values in subsequent model fitting. The same $\lambda$ value was used in fitting the ESD model.

The significance of main and interaction effects was assessed using a permutation test. In each of 1000 permutations, the phenotypes of F2 individuals were randomly reassigned, and the main effect and ESD models were refitted to the permuted data. The improvement in residual deviance resulting from each model fit was calculated and used to generate a null distribution. The residual deviance resulting from each model fit to the observed data was then placed on the tail of the null distribution to determine an approximate *P*-value. The same $\lambda$ value was used in the permutations as with the data.

The results are shown in Table S2. Significant variation in ESD was detected in the x-coordinates of head shape (LDx) in both Paxton and Priest lakes, and in the LDxy trait in Priest Lake. The magnitude of reduction in deviance associated with the ESD model was about one third to one half of that associated with the main effects, although represented only 2-8% of the baseline model deviance. The ESD models identified genotype predictors on chrXI in the Paxton Lake cross and on chrIV, XIII, XVI, and XVII in the Priest Lake cross (not shown).

### Supplementary Tables

**Table S1**. Populations and lakes sampled.

| Lake | Ecotype | Sympatry Type | Females | Males | Latitude | Longitude |
|---|---|---|---|---|---|---|
| Black | Solitary | Solitary | 10 | 10 | 48.77 | -125.10 |
| Bullock | Solitary | Solitary | 9 | 10 | 48.87 | -123.51 |
| Cranby | Solitary | Solitary | 10 | 10 | 49.70 | -124.51 |
| Hoggan | Solitary | Solitary | 7 | 11 | 49.15 | -123.83 |
| Kirk | Solitary | Solitary | 9 | 10 | 49.74 | -124.59 |
| Klein | Solitary | Solitary | 9 | 9 | 49.73 | -123.97 |
| Maxwell | Solitary | Solitary | 9 | 10 | 48.82 | -123.54 |
| Mike | Solitary | Solitary | 10 | 10 | 49.27 | -122.54 |
| Trout | Solitary | Solitary | 8 | 8 | 49.51 | -123.88 |
| Weston | Solitary | Solitary | 9 | 7 | 48.78 | -123.42 |
| Little Quarry | Benthic | Stickleback-sympatric | 11 | 8 | 49.66 | -124.11 |
| Paxton | Benthic | Stickleback-sympatric | 11 | 10 | 49.71 | -124.53 |
| Priest | Benthic | Stickleback-sympatric | 10 | 10 | 49.74 | -124.56 |
| Little Quarry | Limnetic | Stickleback-sympatric | 10 | 10 | 49.66 | -124.11 |
| Paxton | Limnetic | Stickleback-sympatric | 15 | 10 | 49.71 | -124.53 |
| Priest | Limnetic | Stickleback-sympatric | 10 | 13 | 49.74 | -124.56 |
| Blackwater | Sculpin-sympatric | Sculpin-sympatric | 10 | 8 | 50.17 | -125.59 |
| McNair | Sculpin-sympatric | Sculpin-sympatric | 6 | 12 | 50.23 | -125.58 |
| Merrill | Sculpin-sympatric | Sculpin-sympatric | 7 | 8 | 50.06 | -125.56 |
| Morton | Sculpin-sympatric | Sculpin-sympatric | 9 | 11 | 50.12 | -125.48 |
| North | Sculpin-sympatric | Sculpin-sympatric | 7 | 12 | 49.75 | -123.97 |
| Ormond | Sculpin-sympatric | Sculpin-sympatric | 9 | 9 | 50.18 | -125.53 |
| Pachena | Sculpin-sympatric | Sculpin-sympatric | 8 | 10 | 48.84 | -125.03 |
| Paq | Sculpin-sympatric | Sculpin-sympatric | 10 | 11 | 49.61 | -124.02 |
| Roselle | Sculpin-sympatric | Sculpin-sympatric | 9 | 12 | 50.52 | -126.99 |
| Cowichan River | Marine | | 5 | 7 | 48.76 | -123.65 |
| Little Campbell River | Marine | | 9 | 9 | 49.01 | -122.78 |
| Oyster Lagoon | Marine | | 6 | 6 | 49.64 | -123.99 |
| Sayward | Marine | | 7 | 8 | 50.38 | -125.95 |

**Table S2**. Evidence of genetic variation for ecological sexual dimorphism (ESD) in head shape between Limnetics and Benthics. Residual deviance from fitting three models to the data are shown for each lake. The first included only sex and the family covariate and represents the baseline (covariate model). The second added the marker data as main effects (main effects model). The third model (ESD model) also included interactions between sex and genotype, detecting the presence of autosomal genetic variation that has a greater effect in one sex than the other, representing a genetic basis for ESD. The reduction in deviance for each model compared to the preceding model is given in parentheses. Significance levels for improvement in fit of the second and third models are based on permutation test (*$P \leq 0.05$, **$P \leq 0.01$).

| | | **Residual deviance (reduction)** | |
|---|---|---|---|
| **Trait** | | **Paxton Lake** | **Priest Lake** |
| LDx | Covariates model | 303.2 | 257.3 |
| LDx | Main effects model | 289.5 (13.6)* | 208.7 (48.5)** |
| LDx | ESD model | 282.5 (7.0)* | 191.4 (17.3)** |
| LDxy | Covariates model | 325.0 | 250.2 |
| LDxy | Main effects model | 274.0 (51.0)** | 209.0 (41.2) |
| LDxy | ESD model | 268.3 (5.6) | 189.6 (19.4)** |

**Supplementary Figures**

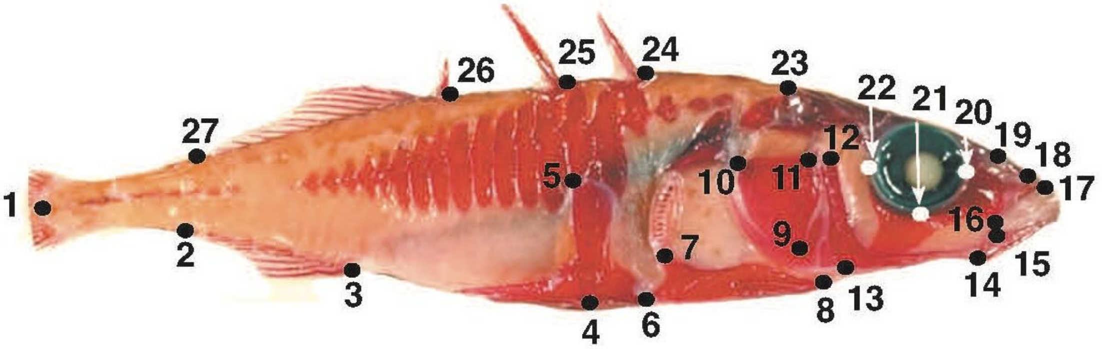


**Figure S1.** Albert et al (2008) landmarks. The same landmarks were used here except the two pelvic girdle landmarks (4 & 5) because the girdle was absent in most Benthics from Paxton and Little Quarry lakes. See Table 1 in Albert et al (2008) for a description. Reproduced from Albert et al (2008) with permission.

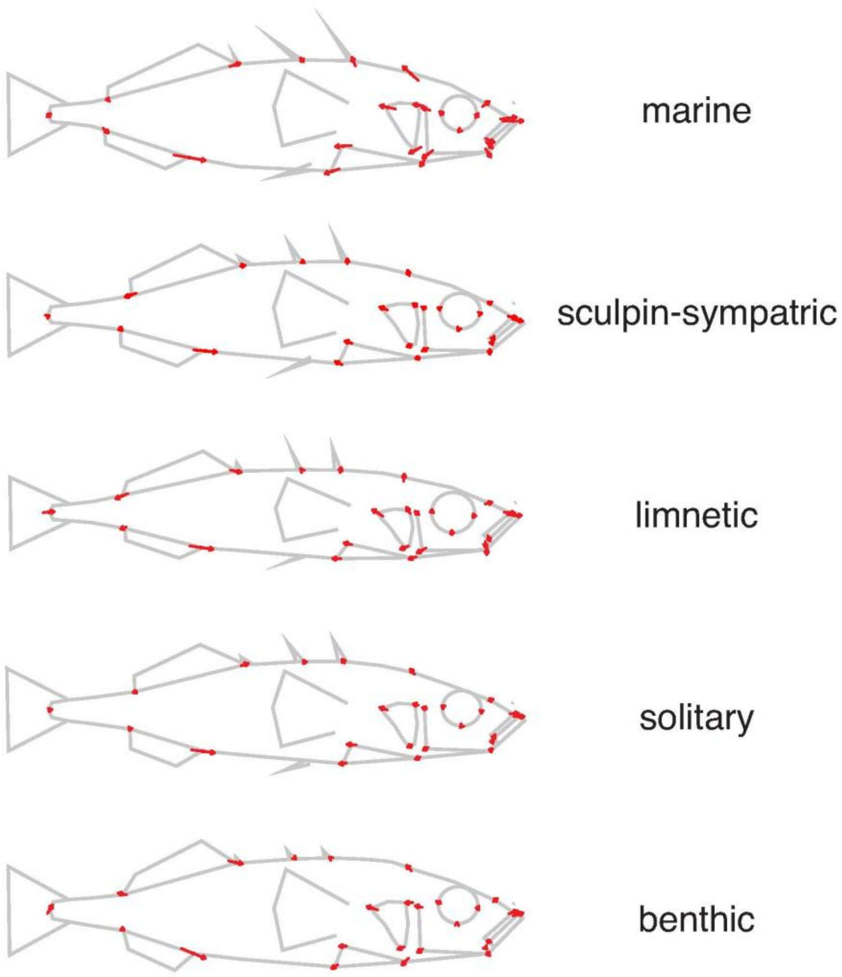


**Figure S2.** Average differences in body shape between males and females of each ecotype. Vectors indicate the difference between average male and average female landmark positions, magnified 3x for visualization.

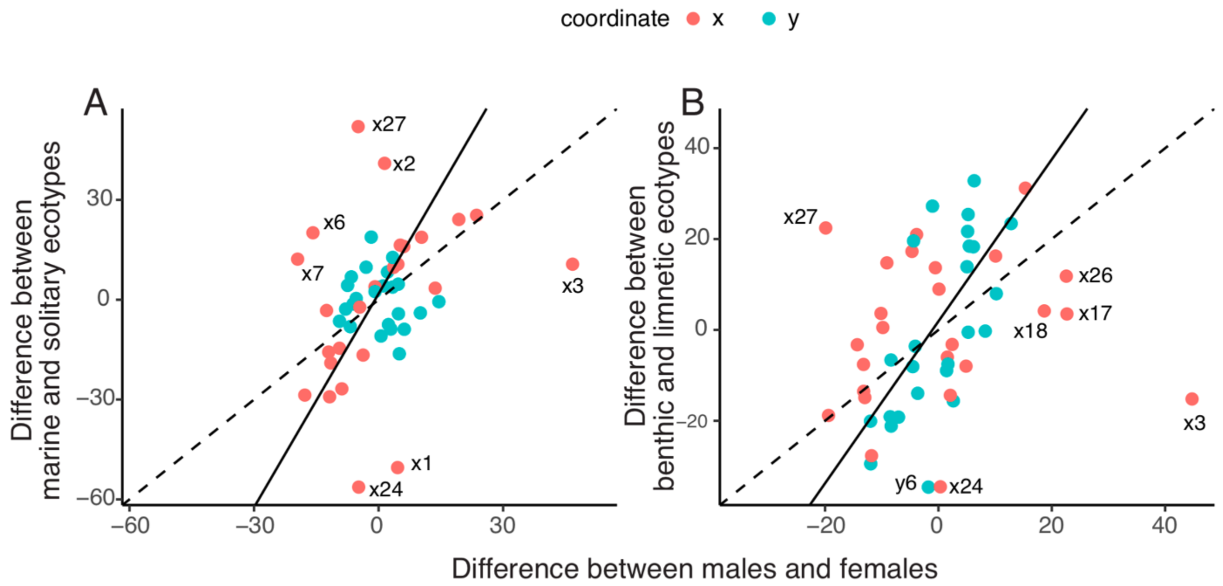


**Figure S3**. Comparison between mean sex and ecotype differences in *x*- and *y*- landmark coordinates. A) marine vs solitary ecotypes. B) Benthic vs Limnetic ecotypes. Values were obtained by fitting a linear mixed model to each landmark coordinate separately in (A) and (B), with sex and ecotype as fixed effects and population as random effect using *blmer*() in the *blme* package (Chung et al. 2013). Means for sex and ecotype were obtained using *emmeans*() (Lenth et al. 2021) and used to calculate ecotype differences and sex differences (males minus females). Dashed lines indicate a 1:1 relationship. Solid lines are reduced major axis regressions calculated using the *sma*() function of the *smatr* package (Warton et al., 2012), leaving out the largest outlying landmark *x*3. Outlier landmark coordinates are indicated.

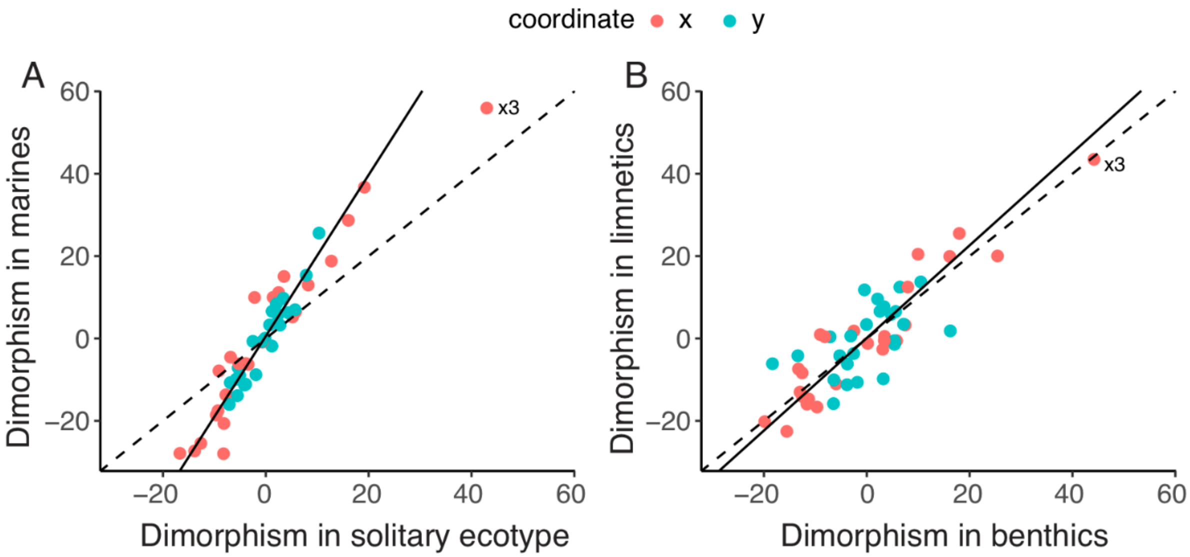


**Figure S4**. Comparison between ecotypes in mean sexual dimorphism in *x*- and *y*- landmark coordinates. A) marine vs solitary ecotypes. B) Benthic vs Limnetic ecotypes. Values were obtained by calculating landmark coordinate means for each sex in every population and then averaging values among populations within each ecotype for males and females separately. Average dimorphism in each ecotype was then calculated as the difference between the male and female means. Dashed lines indicate a 1:1 relationship. Solid lines are reduced major axis regressions calculated using the *sma*() function of the *smatr* package (Warton et al. 2012), leaving out the largest outlying landmark *x*3.

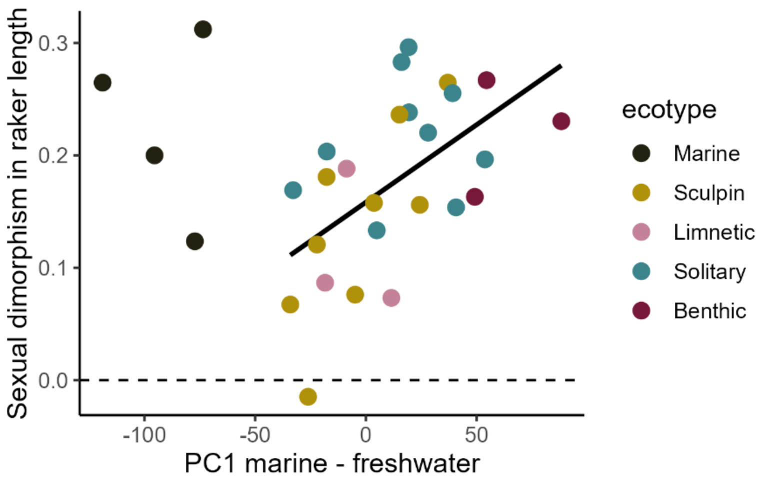


**Figure S5.** Sexual dimorphism in gill raker length in relation to population mean position on the principal marine – freshwater phenotypic axis ($PC1_{M\text{-}FW}$). Gill raker lengths represent the PC1 estimated from size-adjusted gill raker lengths. Each point represents a single population, with the difference between female and male model-fitted means shown along the y-axes.

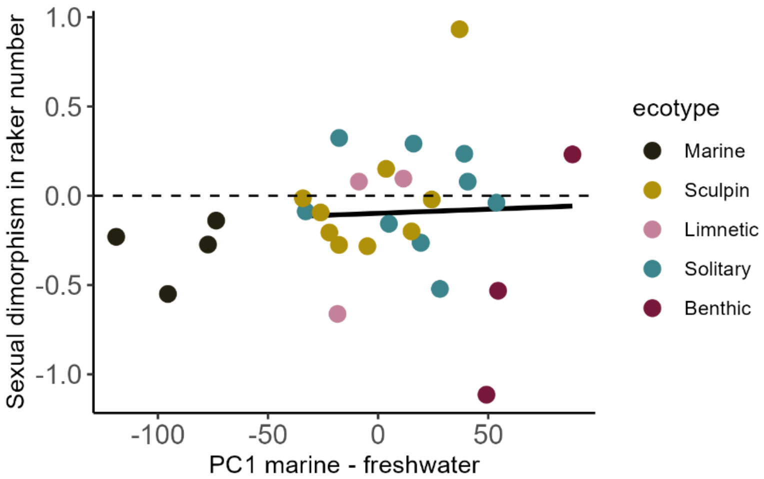


**Figure S6.** Sexual dimorphism in gill raker number in relation to population mean position on the principal marine – freshwater phenotypic axis ($PC1_{M-FW}$). Gill raker numbers are a total count of the upper and lower gill rakers. Each point represents a single population, with the difference between female and male model-fitted means shown along the y-axes.

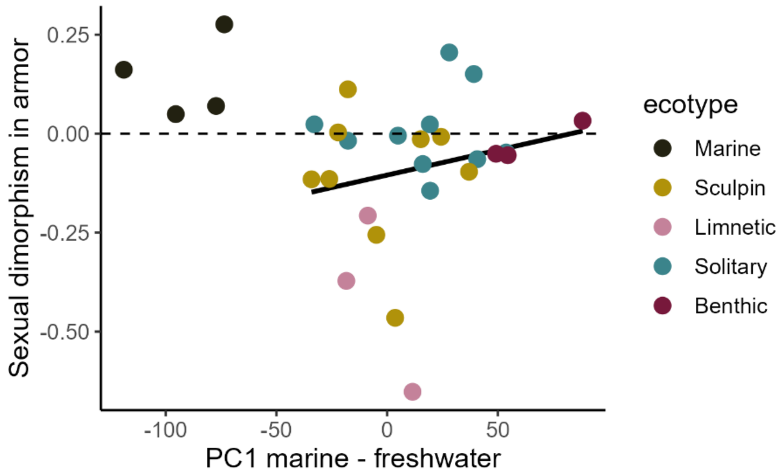


**Figure S7.** Sexual dimorphism in armor traits in relation to population mean position on the principal marine – freshwater phenotypic axis ($PC1_{M\text{-}FW}$). Armor represents the PC1 estimated from size-adjusted armor traits (spine lengths, pelvic girdle, lateral plate number). Each point represents a single population, with the difference between female and male model-fitted means shown along the y-axes.

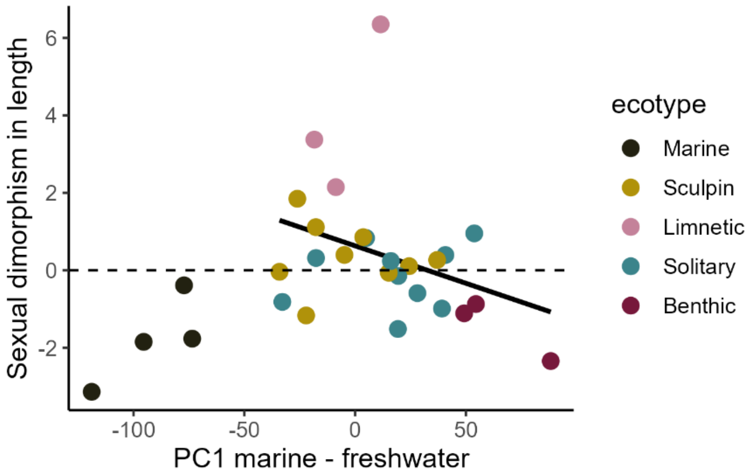


**Figure S8.** Sexual dimorphism in body size (standard length) in relation to population mean position on the principal marine – freshwater phenotypic axis ($PC1_{M\text{-}FW}$). Each point represents a single population, with the difference between female and male model-fitted means shown along the y-axes.

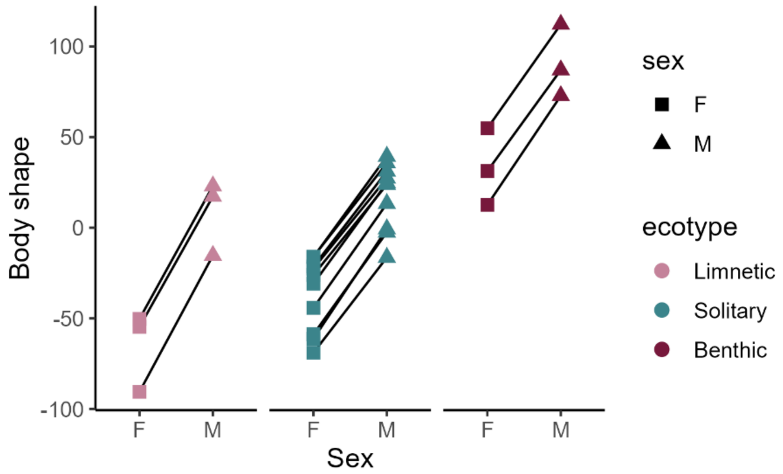


**Figure S9.** Variation in sexual dimorphism in body shape along the limnetic-benthic axis. Each point represents the model-predicted mean body shape $PC1_{Lim\text{-}Ben}$ for males (M) and females (F). Lines connect estimates from the same population.

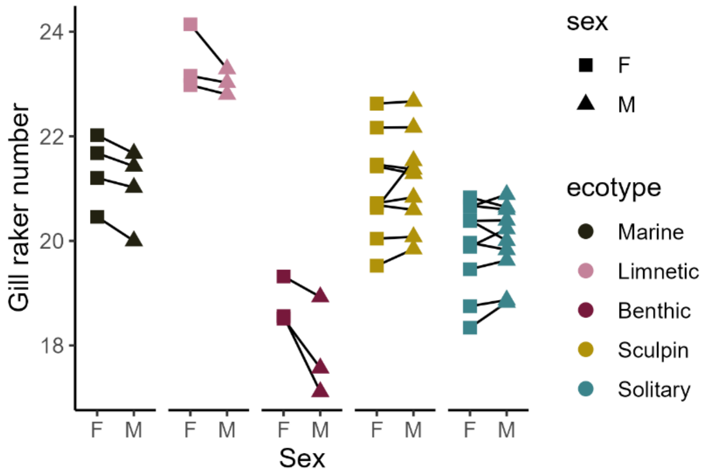


**Figure S10.** Variation in sexual dimorphism in gill raker number. Each point represents the model-predicted mean total gill raker number (upper and lower) for males (M) and females (F), with lines connecting estimates from the same population.

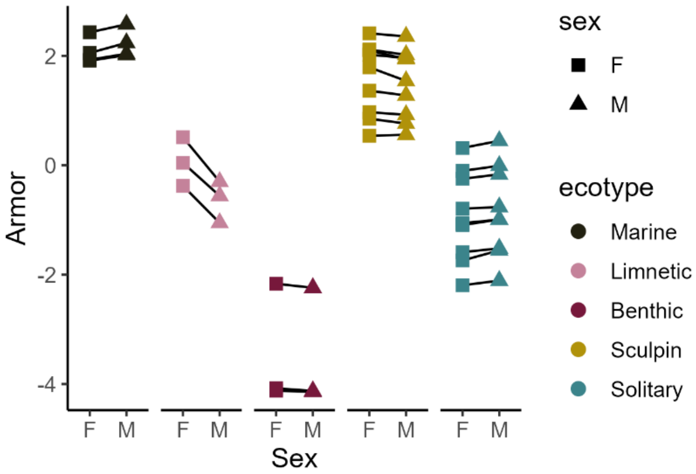


**Figure S11.** Variation in sexual dimorphism in armor traits. Each point represents the model-predicted mean PC1 of size-adjusted armor traits (spine lengths, pelvic girdle, lateral plate number) for males (M) and females (F). Lines connect estimates from the same population.

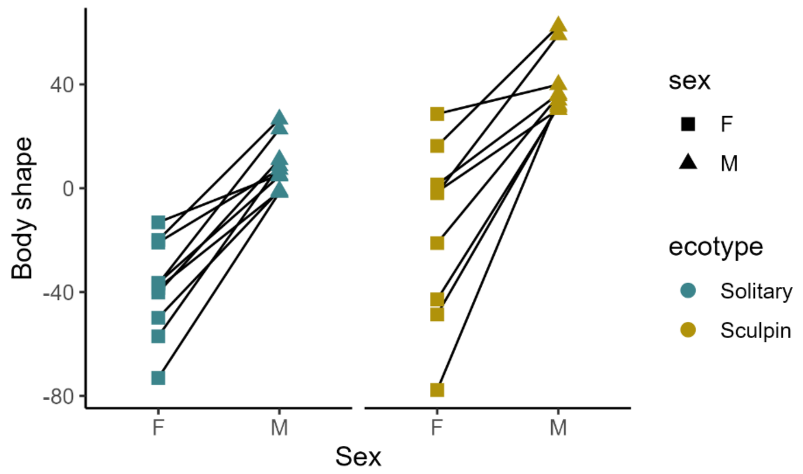


**Figure S12.** Variation in sexual dimorphism in body shape along the principal solitary – sculpin-sympatric axis. Each point represents the model-predicted mean body shape $PC1_{Sol\text{-}Sym}$ for males (M) and females (F). Lines connect estimates from the same population.

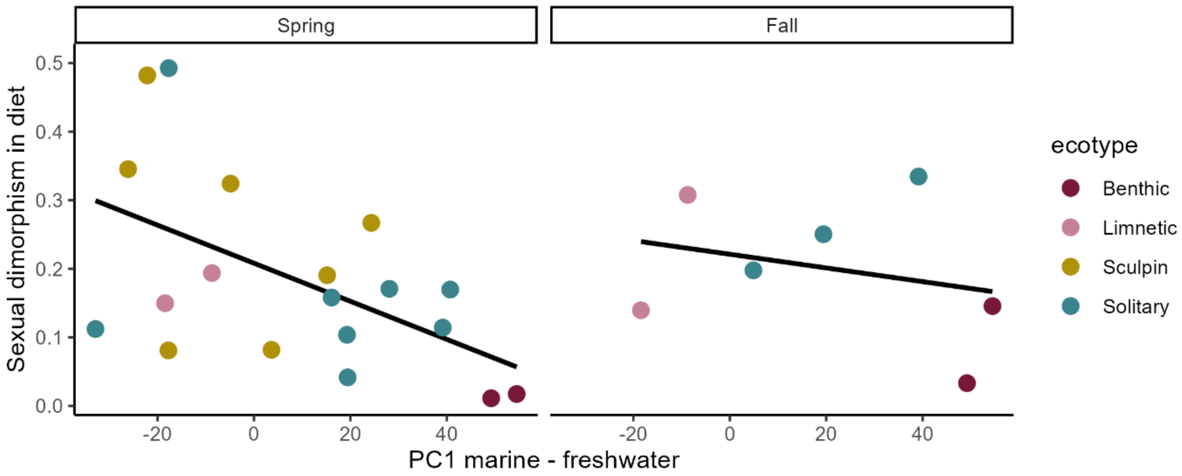


**Figure S13.** Relationship between sexual dimorphism in diet and population mean body shape along the principal marine – freshwater axis ($PC1_{M\text{-}FW}$). Each point represents one population, with dimorphism in diet, quantified as 1 – *PS* (proportional similarity), on the *x*-axis. Specimens were sampled between April and June (left panel) or September (right panel).

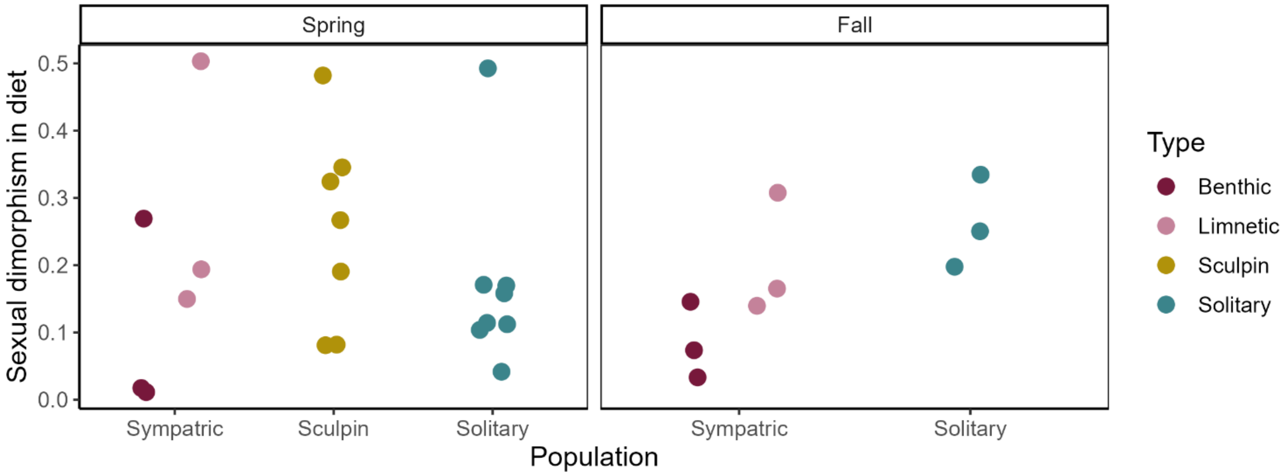


**Figure S14.** Sexual dimorphism in diet in solitary stickleback populations and populations sympatric with another stickleback species or with prickly sculpin. Each point represents the proportional similarity in diet between the sexes. Prey were categorized as “littoral”, “pelagic”, or “other”. Specimens were sampled between April and June (left panel) or September (right panel).
